# Rapid and Sensitive Detection of *Fusarium oxysporum* f. sp. *cubense* Tropical Race 4 Using a RPA-DETECTR Assay

**DOI:** 10.1101/2024.08.15.608054

**Authors:** Jelli Venkatesh, Joanna Jankowicz-Cieslak, Hassan Mduma, Mirta Matijevic, Adel Ali, Isabel Cristina Calle Balbin, Mauricio Soto-Suárez, Cinthya Zorrilla, Pooja Bhatnagar- Mathur

## Abstract

Early and prompt detection of banana wilt pathogen *Fusarium oxysporum* f. sp. *cubense* (Foc), tropical race 4 (Foc TR4), causing global banana crop losses is crucial to curtail the disease spread, minimize damage and implement quarantine measures. We report the first successfully developed DNA Endonuclease Targeted CRISPR Trans Reporter (DETECTR) assay for the highly sensitive and specific detection of Foc TR4, validated across several isolates. Notably, specific crRNA spacer sequences and recombinant polymerase amplification (RPA) primers designed and evaluated for the DETECTR assay exhibited exceptional sensitivity, detecting Foc TR4 with genomic DNA as low as 0.005 ng, and demonstrated a very high specificity, and reproducibility. The RPA-DETECTR assay was validated using a diverse panel of samples, including both target and non-target pathogens confirming its robustness and reliability across different types of samples, ensuring practical application under varied conditions. In terms of throughput, the RPA-DETECTR assays enabled faster detection of Foc TR4 than other molecular diagnostics including PCR, quantitative PCR (qPCR) and droplet digital PCR (ddPCR) analyses, making this assay suitable for point-of-care (POC) detection of Foc TR4, and facilitating rapid decision-making and immediate response. This rapid detection capability is crucial in restricting any potential transboundary movement as part of ongoing disease management efforts.

## Introduction

*Fusarium oxysporum* f. sp. *cubense* (Foc) is a fungal pathogen responsible for Fusarium wilt, a significant disease affecting bananas and plantains with a history of impact on global banana cultivation (Were et al. 2023). In the first half of the last century, Fusarium wilt caused by Foc race 1 (Foc R1) characterized as one of the most destructive plant diseases ever documented led to substantial losses in Gros Michel banana plantations in Latin America (Moore et al. 1995). Despite extensive efforts, disease management failed, and production recovery relied on replacing Gros Michel with resistant varieties from the Cavendish subgroup, subsequently becoming the predominant variety for the commercial banana production (Ploetz 2015; Ploetz et al. 1994). Until 2018, over 50% of global banana production depended on this single variety (Lescot 2020). However, commercial banana production now faces a renewed threat from another Foc race, tropical race 4 (Foc TR4), belonging to vegetative compatibility group (VCG) 01213/16 (EFSA PLH Panel et al. 2022) which is highly pathogenic to the Cavendish subgroup with currently no viable solutions for eradication or effective management. The Foc TR4, genetically reclassified as *F. odoratissimum* (Maryani et al. 2019), though this classification is not widely accepted by the phytopathology community.

The lack of alternative, highly productive, and market-acceptable resistant banana varieties accentuates Foc TR4 threat (Ploetz 2015). Additionally, the susceptibility of local banana varieties, including plantains and cooking banana to Foc TR4 exacerbates the threat not only to economic revenue but also food security (Ploetz 2006; Zhan et al. 2022). The persistent expansion of Fusarium wilt (caused by Foc TR4) adversely impacts the food security of small-scale farmers who depend on local banana varieties as a primary food staple (van Westerhoven et al. 2022). Management of the disease is extremely difficult as currently there are no efficient strategies for the prevention and control of Foc TR4 (Ploetz 2006; Bubici et al. 2019; Pegg et al. 2019). The available means of effective control measures are presently limited to exclusion and containment strategies. Nonetheless, to facilitate successful control measures, it is crucial to have accurate diagnostic tools for early detection. The longer the Foc TR4 goes unnoticed in a specific area, the greater the risk of its further spread, posing an escalating threat to the multi-billion-dollar banana industry. Additionally, the livelihoods and food security of millions of people worldwide, who depend on bananas and plantains, are increasingly jeopardized as a consequence (FAO 2022; Ordonez et al. 2015).

An immediate and critical priority is developing diagnostic tools that are highly efficient, sensitive, and specific. Ideally, these would offer an efficient early warning in the context of crop production, thereby substantially mitigating the impact of Foc TR4 outbreaks and minimizing associated economic losses. Several nucleic acid based methods including PCR, quantitative PCR (qPCR) and droplet digital PCR (ddPCR) have been employed for the detection of Foc TR4 (Lin et al. 2009; Dita et al. 2010; Li et al. 2013a; Li et al. 2013b; Lin et al. 2013; Peng et al. 2014; Yang et al. 2015; Lin et al. 2016; Carvalhais et al. 2019; Thangavelu et al. 2019; Matthews et al. 2020; Aguayo et al. 2017; Ordóñez et al. 2019; Lovera et al. 2024). Generally, these methods target genomic regions associated with SCAR and RAPD markers, as well as other targets like genomic regions coding for *Secreted in Xylem* (*SIX*) genes and hypothetical proteins, to achieve accurate and reliable detection. Despite its widespread use, these approaches have limitations, particularly in resource-constrained settings due to the necessity for large and expensive thermal cyclers. A promising alternative has emerged in recent years with advent of loop-mediated isothermal amplification (LAMP) for rapid TR4 detection, which allowed rapid and efficient amplification at a constant temperature without the need for thermocycling (Li et al. 2013a; Ordóñez et al. 2019). However, addressing the challenge of non-specific amplification remains a significant obstacle to optimizing accurate LAMP assays (Rolando et al. 2020). Ideal diagnostic methods should enable on-site testing where infected plants are located, known as point-of-care (POC) testing. These methods should utilize instruments that are portable, cost-effective, exhibit high specificity for the target molecule, and provide rapid results (LauandBotella 2017).

More recently, the Clustered Regularly Interspaced Short Palindromic Repeats (CRISPR)-Cas has rapidly evolved over the past few years for precise genome editing applications as well as detection of nucleic acids (Kaminski et al. 2021) offering tremendous potential for the detection of nucleic acids that meet the criteria of sensitivity, specificity, affordability, speed, user-friendliness, and minimal equipment requirements (Huang et al. 2023). The CRISPR/Cas12a-based nucleic acid detection system, utilized in the DNA Endonuclease-Targeted CRISPR Trans Reporter (DETECTR) assay, functions by recognizing a specific target DNA sequence with a short T-rich (5′-TTTV-3′) protospacer adjacent motif (PAM). When the Cas12a-crRNA ribonucleoprotein complex binds to and cleaves the target DNA, it triggers trans-cleavage activity. This activity leads to the nonspecific cleavage of single-stranded DNA (ssDNA) reporters/probes. The cleavage of these reporters can be visualized through real-time PCR or lateral flow assay (LFA), providing a rapid and highly sensitive method for detecting the presence of specific genomic targets (Chen et al. 2018; Yamano et al. 2016).

For purposes of diagnostics, pathogen-specific genomic targets can be detected by utilizing suitable crRNAs that guide the detection system, in conjunction with an ssDNA reporter or probe. This approach has been successfully employed in the detection of coronaviruses, including SARS-CoV-2, human papillomavirus (Chen et al. 2018) and certain strains of pathogenic bacteria (Yao et al. 2018; Kang et al. 2021) demonstrating its effectiveness in identifying pathogens with high specificity (Broughton et al. 2020). To enhance the detection sensitivity, CRISPR/Cas12a system have been integrated with nucleic acid amplification methods, such as PCR, LAMP, recombinant polymerase amplification (RPA) (Luo et al. 2021; Wang et al. 2023; Marqués et al. 2022). These combinations enhance sensitivity for detecting target nucleic acids, offering a rapid, robust and efficient identification and analysis method (Luo et al. 2021; Wang et al. 2023; Marqués et al. 2022).

In this study, we developed the DETECTR platform, integrating RPA, CRISPR/Cas12a system and LFA for precise identification of Foc TR4. Specific genomic regions of Foc TR4 were targeted for DETECTR specificity. Rigorous testing validated the integrated platform’s effectiveness in identifying Foc TR4 in infected banana tissue and panels. The use of lateral flow strips enabled straightforward visual interpretation. This advancement promises on-site applications, aiding POC diagnostics and swift decision-making in agriculture. Our findings highlight the potential of the DETECTR system for early and accurate Foc TR4 detection, aiding effective disease management and safeguarding banana crops.

## Materials and methods

### Foc strains and diagnostic panel

Several *Fusarium oxysporum* strains viz. garlic pathogen *F. oxysporum* (Fo), Foc R1, and Foc TR4 diverse collections were included in the present study across experiments (See Additional file 1: Table S1). For validation of DETECTR assay, we used a diverse panel of banana samples from Instituto Colombiano Agropecuario (ICA), Colombia. The panel included DNA from un-inoculated controls and infected Cavendish and Plantain 4 Filos plants, and banana genomic DNA samples spiked with different endophytes, Foc R1, Foc TR4, Foc subtropical race 4 (Foc STR4) and *Ralstonia solanacearum*.

### Inoculum preparation

The solid inoculum was prepared as previously described (Ndayihanzamaso et al. 2020) with slight modifications. Briefly, three grams of Mung bean (*Vigna radiate*) were measured and put in a 500 ml Erlenmeyer flask (Duran) to which 250 ml of sterile distilled water was added, capped with a sterile cotton wool covered with Aluminium foil and autoclaved at 120°C for 15 min. Fungal hyphae were transferred from a long-term freeze storage vial onto potato dextrose agar (PDA) plates and incubated at room temperature (25°C) for a period of 5-7 days. Subsequently, 2-3 Foc mycelial plugs (∼5 mm in diameter), were then cut from the growing edges of Foc colonies on PDA plates and transferred to bags containing sterile millet seeds, as well as to a a flask containing mung bean solution (Ndayihanzamaso 2020). The suspension was placed in a shaking-incubator and rotated at 150 rpm for 4-5 days at room temperature. Spores were thereafter separated from the mycelia by filtration through a double-layer of sterile cheesecloth, and the spores transferred into 50-ml falcon tubes (Termo Fischer Scientific, Massachusetts, USA). The spore concentration in the suspension was determined using a haemocytometer and adjusted to a final concentration of 1×10^6^ conidia ml^−1^ and immediately used for inoculations. Spore viability was confirmed by plating onto PDA media.

### Banana Foc TR4 inoculations

*In vitro* raised Cavendish banana plantlets were transplanted in a plastic pot (10 × 10 × 12 cm) filled with potting mixture containing 1:1 soil and peat (K-SUBSTRATES select, Klasmann-Deilmann, Geeste, Germany) under contained greenhouse environment at Seibersdorf Laboratories in Lower Austria. Banana plants were inoculated as mentioned above at four-leaf stage and maintained at an average temperature of 24±2°C and a 16-h photoperiod and were watered according to requirements.

### Genomic DNA extraction

Genomic DNA from Foc mycelia was extracted using the DNeasy Plant Mini Kit (QIAGEN, Hilden, Germany). Genomic DNA from plant samples were extracted using the DNeasy Plant Mini Kit (QIAGEN, Hilden, Germany). Four weeks post-inoculation, plants were delicately extracted from their pots and rinsed under tap water. Subsequently, pseudostems and corms were dissected to inspect for vascular discoloration, and samples were collected and used for genomic DNA was extraction. The concentration of DNA was measured using a NanoDrop One spectrophotometer (Thermo Fisher, Waltham, MA, USA).

### Target selection for the DETECTR assay, crRNA design and synthesis

Previously validated Foc TR4 specific sequence, SeqA (Ordóñez et al. 2019) and a sequence characterized amplified region (SCAR) marker (Li et al. 2013a), were selected for DETECTR based diagnostic markers development. CRISPR/Cas12 target specific primers were designed as per the recommendations of the TwistAmp^TM^ Liquid Basic kit (TwistDx, Cambridge, UK). Specific RPA primers and crRNA spacer for the RPA-DETECTR assays (Table 1) were designed and commercially produced at Eurofins (Ebersberg, Germany). Lba Cas12a crRNA spacer sequence was selected based on PAM (TTTV) within the Foc TR4 specific DETECTR target region. For target specific crRNA synthesis, DNA oligo containing crRNA spacer, crRNA scaffold and T7 promoter sequences were synthesized by Eurofins (Ebersberg, Germany). Oligos were used for the double-stranded DNA (dsDNA) synthesis using Q5 polymerase (New England Biolabs, Frankfurt am Main, Germany) in a reaction consisting of 25 µl Q5 High-Fidelity 2× Master, 2.5 µl of forward (10 µM) and 2.5 µl of T7-R reverse (10 µM) primers and 1 µL crRNA-DNA oligo (10 µM) and distilled water to make up the final volume (50 µl). The PCR cycling conditions were 98°C for 30 s, 38 cycles of 98°C for 5 s, 56°C for 10 s and 72°C for 15 s and a final extension of 72°C for 2 min. Amplified crRNA dsDNA was gel purified and eluted and subsequently used for *in vitro* crRNA synthesis following the HiScribe T7 High Yield RNA Synthesis Kit (New England Biolabs, Germany). CrRNA was synthesized from dsDNA template by overnight incubation at 37°C using HiScribe T7 High Yield RNA Synthesis Kit (New England Biolabs, Frankfurt am Main, Germany). Transcribed crRNA was cleaned up using Monarch^@^ RNA Cleanup Kit (New England Biolabs, Frankfurt am Main, Germany), crRNA quality and quantity was analysed by gel electrophoresis and Nanodrop, and stored at −80°C for subsequent studies.

**Table 1.**
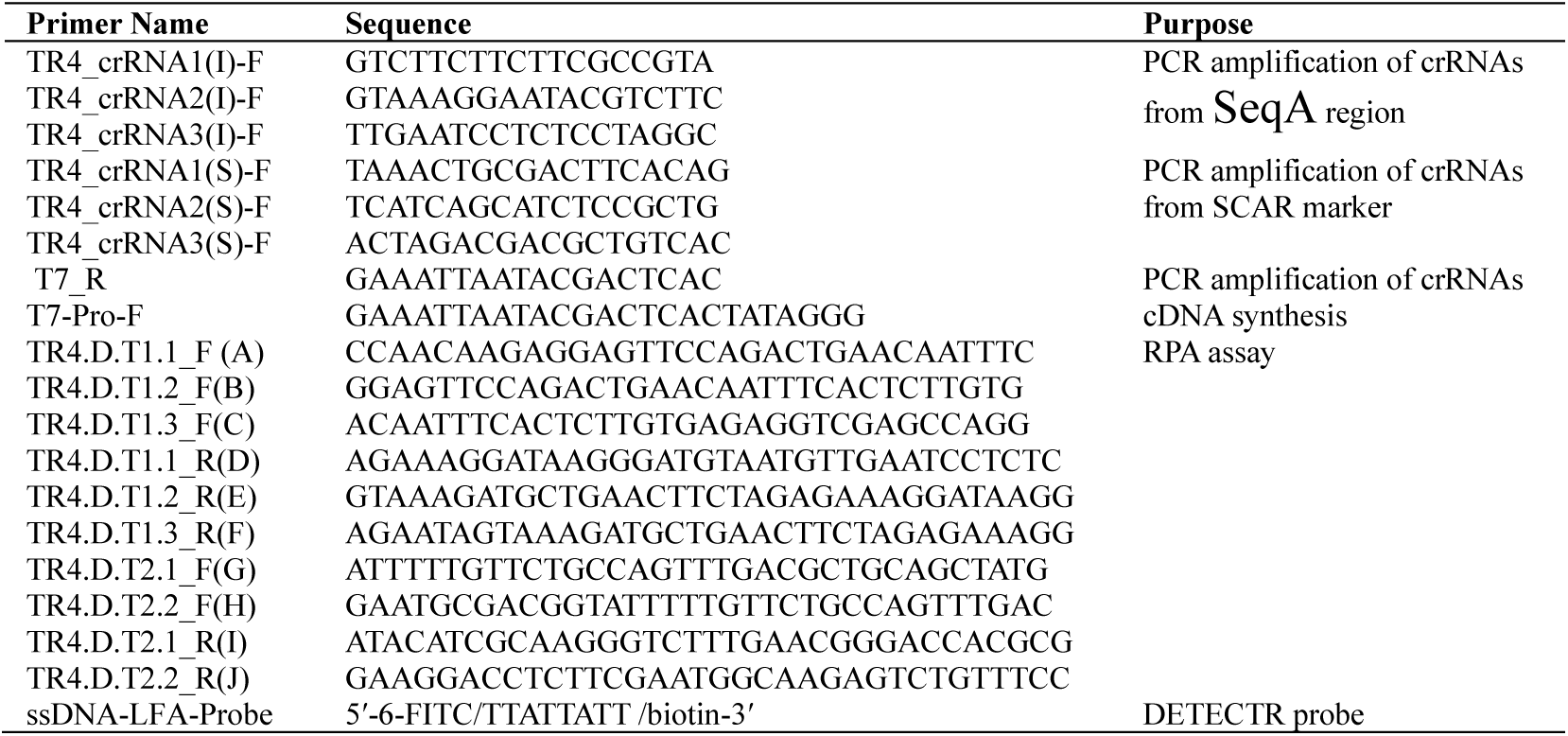
List of primers used in the present study.

**Table 2.**
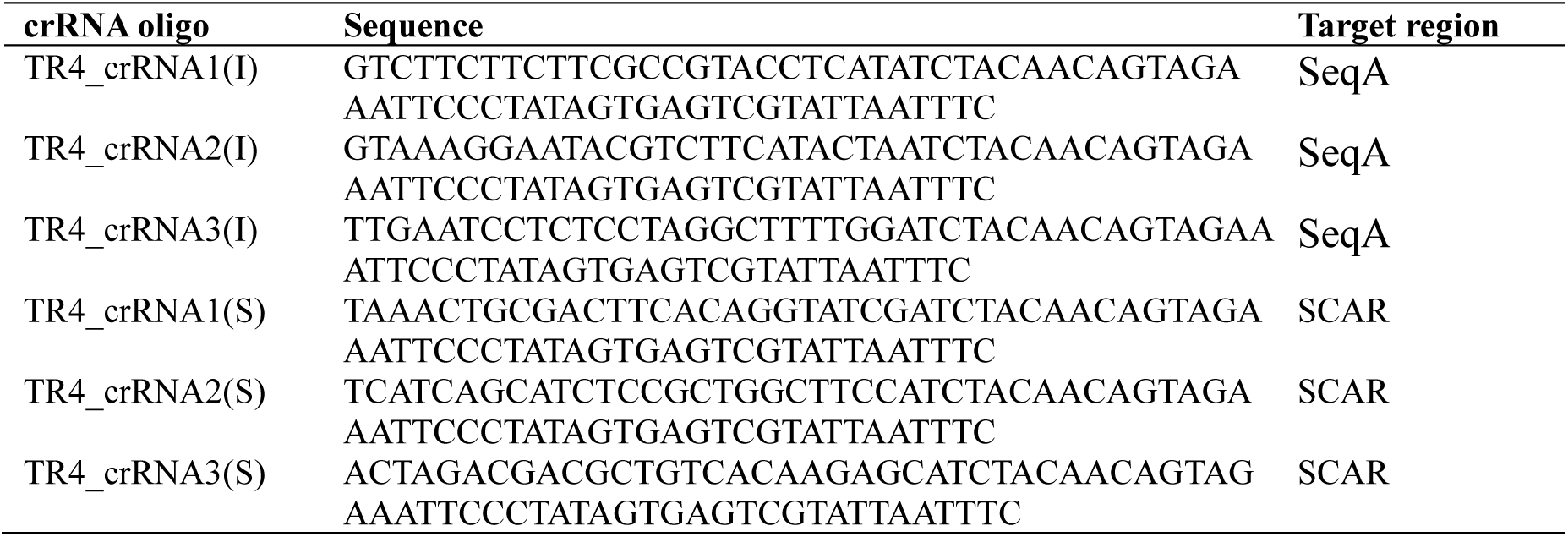
DETECTR crRNA oligos used for amplification.

### Validation of Cas12a cleavage activity with different crRNAs

PCR amplification of DETECTR targets were carried out in a reaction volume of 25 μl using Foc TR4 specific primer pair listed in the table (**Table 1**), The reaction included 2.5 μl 10× Ex Taq buffer, 0.5 μl of 10 μM of forward and reverse primers, 0.5 μl of 50 ng DNA template, and 0.25 μl Ex Taq DNA polymerase Hot Start Version (Takara Bio, Tokyo, Japan) with total volume make up with addition of RNase-free water. PCR cycle conditions were as: 95°C for 3 min; 38 cycles of 95°C for 20 s, 55°C for 20 s, 72°C for 20 s; 72°C for 3 min. The resulting PCR amplicons were analysed by 2.0% agarose gel electrophoresis.

Cleavage activity was performed in a reaction mixture containing 3 μl of 10× NEBuffer r2.1, 300 nM crRNA, and 30 nM EnGen Lba Cas12a (New England Biolabs, Germany) with final volume 27 μl made up with RNase-free water. Reaction mixture was preincubated at 25°C for 10 min and then 3.0 μl of PCR product was added and incubated for 30 min at 37°C, then 1 μl of Proteinase K (New England Biolabs, Germany) was added to the sample and incubated 10 min at room temperature before samples were analysed by 2.0% gel electrophoresis.

### Verifying Cas12a trans-cleavage activity

To identity the suitable crRNAs with an efficient Lba Cas12a trans-cleavage activity, a nucleotide reporter consisting of fluorescein isothiocyanate (FITC) fluorophore and biotin tags (5′FITC-TTATTATT-3′biotin) was synthesized by Eurofins (Ebersberg, Germany) was used in the trans-cleavage assay. Trans-cleavage assay was performed in a reaction volume of 30 μl containing 1× NEBuffer r2.1, 30 nM reporter, 300 nM crRNA, and 100 nM Lba Cas12a and 3 μl Foc strain specific PCR product and the reaction mixture was incubated in thermal cycler (C1000 Touch^TM^, Bio Rad) for 30 min at 37°C. For LFA detection, 100 μl of HybriDetect assay buffer (Milenia Biotec, Germany) was added to the reaction and allowed to incubate at room temperature for 2 min. A lateral flow strip (Milenia Biotec, Germany) was immersed in the reaction for 2 min and observed for the development of a visible color in both the test and control lines.

### Primer screening for the RPA assay

RPA was performed as per the recommended protocol of TwistAmp^TM^ Liquid Basic (TwistDx, Cambridge, UK) with some modifications. Briefly, reaction mix containing 12.5 μl 2× reaction buffer, 1.125 μl of each of 10 μM forward and reverse primers, 4.0 μl of dNTPs mix (10 mM), and 2.5 μl 10× Basic E-mix, and 0.25 μl sterile water was prepared and vortexed and spin briefly, following the addition of 1.25 μl 20× core reaction mix and subsequently mixed by 10× full inversions and brief spin. Thereafter, 1.0 μl template and 1.25 μl of 280 mM Magnesium acetate (MgOAc) were added to the PCR tubes containing 22.75 μl of reaction mix. The mixture was then thoroughly mixed well (6×) and incubated at 37°C for 25 min in a thermal cycler (C1000 Touch^TM^, Bio Rad). The resulting product was analysed by 2.0% agarose gel electrophoresis after PCR clean up using QIAquick PCR Purification Kit (QIAGEN, Hilden, Germany).

## DETECTR assays

### Validation of crRNAs

The variability of DETECTR system with different crRNAs were analysed through the utilization of a LFA system. RPA reactions were performed under the optimized conditions as described in TwistAmp^TM^ Liquid Basic kit (TwistDx, Cambridge, UK). To amplify the target sequences, a reaction comprising of 12.5 μl of rehydration buffer (TwistDx, Cambridge, UK), 1.125 μl each of forward and reverse primers (10 μM), 4 μl of dNTPs mix (10 mM), and 2.5 μl 10× Basic E-mix, was adjusted to a final volume of 22 μl with double-distilled water, and vortexed briefly. This mixture was then combined with 1.25 μl of 20× core reaction mix, 1.0 μl of template DNA (5 ng), and 1.25 μl MgOAc (280 mM). The amplification reaction was incubated at 39°C for 25 min. Following incubation, 2 μl of RPA product was added to the reaction mix consisting of 1 μl of Lba Cas12a (1 μM), 1 μl of crRNA (100 ng), 1 μl of ssDNA-reporter (10 μM) and 15 μl of double-distilled water and the reaction mix was incubated at 37℃ for 25 min. For the LFA assay, 100 μl of HybriDetect assay buffer was added to each tube containing 20 μl of DETECTR product. A lateral flow strip (Milenia Biotec, Germany) was immersed in the reaction for 2 min, and then observed for the development of a visible color in test (T) and control (C) lines.

### Sensitivity and specificity analysis

The sensitivity of the optimized DETECTR was validated using fungal genomic DNA template, with a 10-fold serial dilution ranging from 5.0 ng to 0.005×10^−3^ ng per reaction. Additionally, the specificity of the assay was confirmed utilizing genomic DNA from various Foc strains (See Additional file 1: Table S1). DETECTR assay was performed as mentioned in the above section. Briefly, 2 μl of RPA product was used for DETECTR assay. The ssDNA reporter labelled with FITC and biotin was employed (**Table 1**). A HybriDetect LFA dip strip (Milenia Biotec, Germany) was carefully placed in the Eppendorf tube containing 100 μl of HybriDetect assay buffer and 20 μl of DETECTR product, and incubated for 2 min at room temperature. A negative control was included for visualization of background signals, utilizing an addition of ssDNA reporter with RNase-free water as the template. Experiments were conducted in triplicate. The intensity of the T-line was quantified using ImageJ software (https://imagej.net/software/imagej/).

### PCR, qPCR and ddPCR analyses

PCR reactions were conducted in a 25 μl volume, including 5 μl of PrimeSTAR GXL buffer, 2 μl of 2.5 mM dNTPs, 0.5 μl of each reverse and forward primer (AF primers), 1 μl of fungal genomic DNA at various concentrations, 0.5 μl of PrimeSTAR GXL Taq DNA polymerase, and 15.5 μl of sterile water. The PCR protocol followed an initial denaturation at 95℃ for 3 min, followed by 34 cycles of 98℃ for 10 s, 62℃ for 15 s, and 68℃ for 20 s, with a final extension at 68℃ for 3 min.

Sensitivity was further assessed using qPCR using AF primers (**Table 1)**, qPCR amplification of DETECTR target sequences were carried out in 20 μl reaction volume using Foc TR4 specific AF primer pair, 10 μl of 2 × Sso Advanced universal SYBR Green Supermix (Bio-Rad) and 1.0 μl of genomic DNA template, 0.3 μl of 10 μM of forward and reverse primers with volume make up (20 μl) with addition of RNase-free water. The qPCR program was followed as initial denaturation at 98°C for 5 min, followed by 40 cycles of denaturation at 95°C for 15 s, annealing and extension at 62°C for 30 s, followed by a melt melting curve analysis (60°C to 95°C: increment 0.5°C per every 5 s). qPCR was conducted on a CFX 96 instrument (Bio-Rad), and sensitivity was verified based on cycle threshold (Ct) values obtained at different DNA sample concentrations and data analysed using Bio-Rad CFX Maestro software 1.1.

The ddPCR was carried out using a QX200™ Droplet Digital™ PCR system (Bio-Rad, USA). The 22 μl ddPCR reaction mixture included 11 μl of QX200 ddPCR EvaGreen Supermix, 0.33 μl of each 10 μM forward and reverse primer (AF primers), 1 μl of fungal genomic DNA, and 9.34 μl of sterile distilled water. 20 μl of this reaction mix was then placed into the sample wells of a DG8™ cartridge (Bio-Rad, Hercules, CA, USA), and 70 μl of droplet generation oil was added to the oil wells (Bio-Rad, Hercules, CA, USA), followed by covering with DG8™ Gaskets (Bio-Rad, USA). The droplets were generated using a QX200™ droplet generator (Bio-Rad, USA). These droplets (40 μl) were then transferred to a 96-well plate (Bio-Rad, Hercules, USA), sealed with Pierceable Foil Heat Seal (Bio-Rad, USA) using a PX1™ PCR plate sealer (Bio-Rad, USA), and subjected to thermal cycling on a C1000 Touch™ Thermal Cycler (Bio-Rad, USA). The thermal cycling conditions were as 95°C for 5 min, 40 cycles of 95°C for 30 s, 60°C for 60 s, with a ramp rate of 2°C/s. The reaction was then subjected to a final extension at 98°C for 10 minutes and held at 4°C. After amplification, the droplets were analysed with a QX200™ Droplet Reader (Bio-Rad, Hercules, USA), and the amplicons were quantified using QX Manager 1.2 Standard Edition software (Bio-Rad, Hercules, USA).

### Statistical analysis

Data analysis was performed with Excel 2019. The experimental data are presented as the means ± SEs.

## Results

### Molecular validation of the Foc TR4 strains

To verify the identity of the Foc TR4 strains, samples from different regions were subjected to PCR validation using the W2987 primers (Li et al. 2013b) and LAMP primers specific to SCAR and SeqA genomic regions (Li et al. 2013a; Ordóñez et al. 2019), which have been previously reported to be specific to Foc TR4. The PCR results demonstrated successful amplification with the Foc TR4 strains, while no amplification was observed with the Foc R1 strains (**Fig. 1**). Consequently, the SCAR and SeqA genomic regions were targeted for the development of the DETECTR assay.

**Fig 1.**
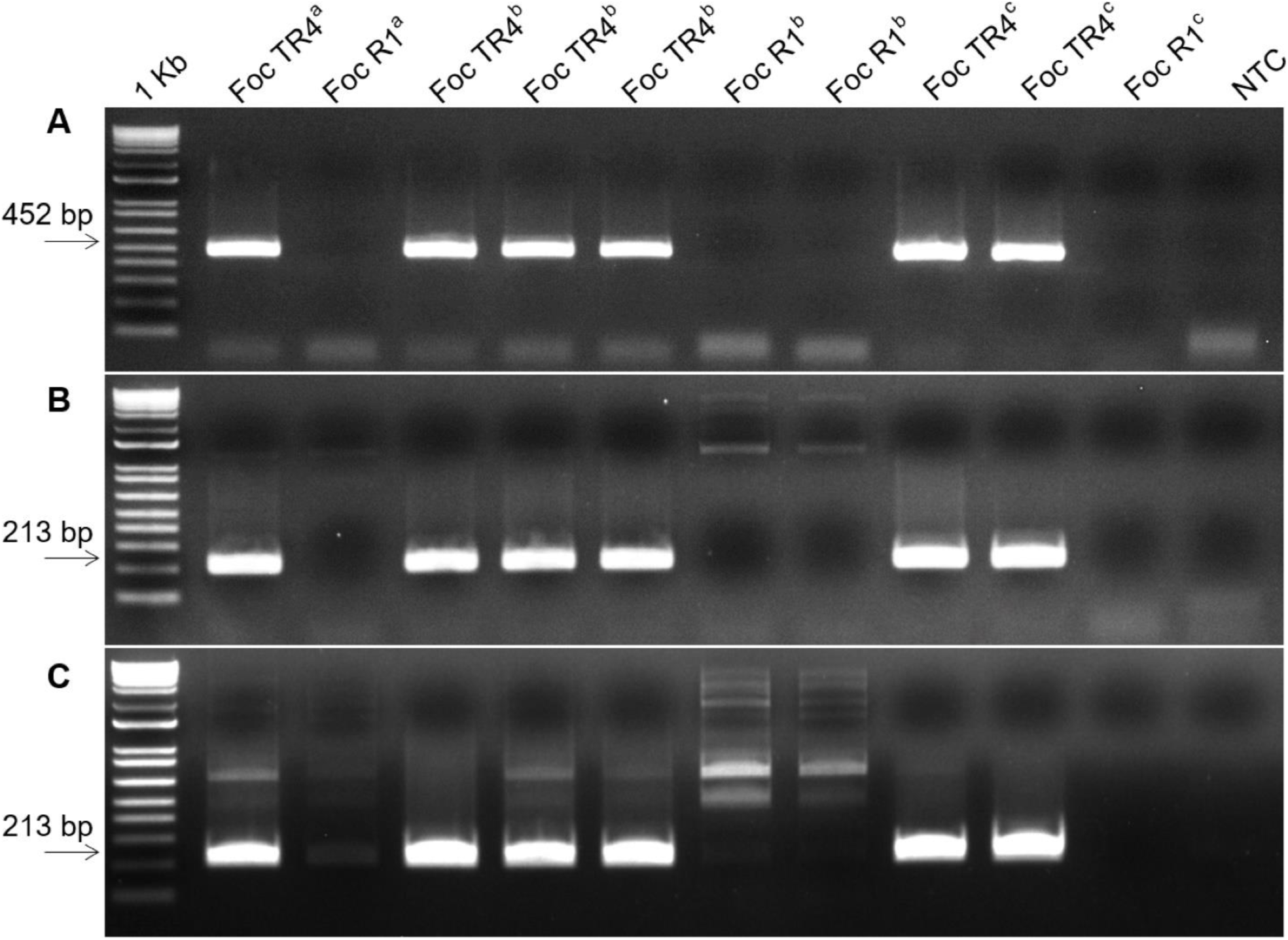
PCR validation of TR4 strains. **A**, PCR was performed using W2987 (Li et al. 2013b). **B**, PCR was performed using SeqA-specific marker (Ordóñez et al. 2019). **C**, PCR performed using SCAR-specific marker (Li et al. 2013a). ^a^ Foc TR4 and Foc R1 from Stellenbosch University, South Africa, ^b^ Foc TR4 strains (Foc TR4 reference, MFocR4T Ref.Agro CP-M 4, MFocR4T Ref.Agro CP-M 5), Foc R1 strains (M C R1 M3723M000199 1, M C R1 M3723M000199 4) from Instituto Colombiano Agropecuario (ICA), Colombia, ^c^ Foc TR4 (190048 and 190098) and Foc R1 from Agrosavia, Colombia. NTC, no template control.

### Identification of suitable crRNA

Three crRNAs were developed and tested for each of the two Foc TR4 specific SCAR marker and SeqA target regions (See Additional file 1: **Fig. S1**), using Foc TR4 VCG 01213/16 strain (**Fig. 2A and D**). The crRNAs designed for these target regions exhibited variable efficiency in terms of their cleavage activities. Notably, all three crRNAs from SCAR region exhibited high cleavage activity, as depicted in **Fig. 2B**, cleaving the target DNA efficiently to produce predicted fragments, suggesting their appropriateness for the assay. However, in contrast only one crRNA (crRNA1(I)) from the SeqA region demonstrated high CRISPR cleavage activity with crRNA2(I) showing a detectable cleavage activity (**Fig. 2E**), suggesting the suitability of two out of three tested crRNAs (crRNA1(I) and crRNA2(I)) from the SeqA region.

**Fig. 2.**
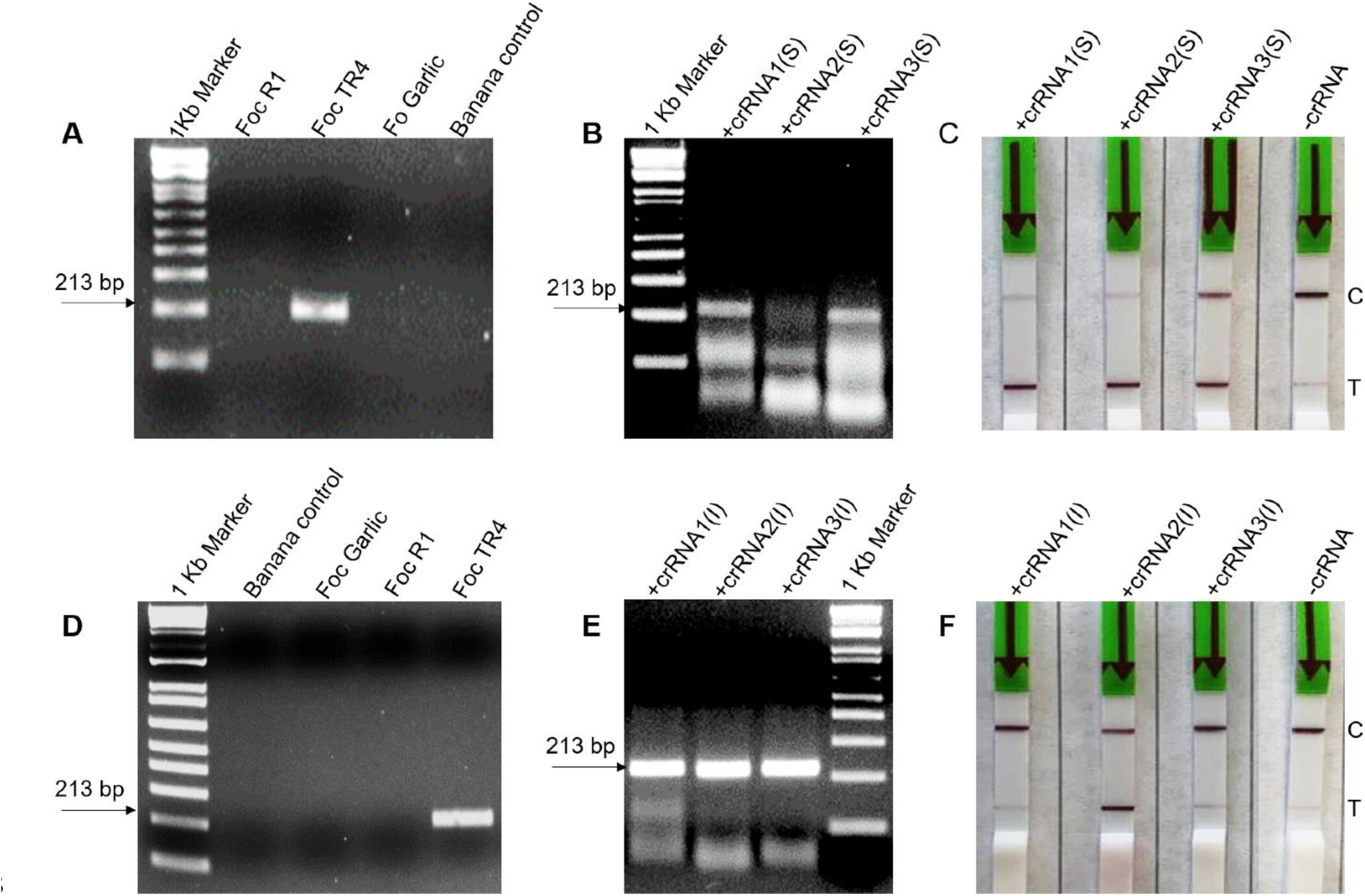
PCR-Based DETECTR Assays for Foc TR4 Detection. **A**, PCR evaluation of the specificity of Foc TR4-specific primers (Li et al. 2013a). **B**, *In vitro* cleavage of Foc TR4-specific PCR products from SCAR region by Lba Cas12a and crRNAs. **C**, Visual representation of lateral flow strips resulting from the SCAR region DETECTR assay. **D**, PCR evaluation of the specificity of Foc TR4-specific primers (Ordóñez et al. 2019). **E**, *In vitro* cleavage of Foc TR4-specific PCR products from SeqA region by Lba Cas12a and crRNAs. **F**, Visual representation of lateral flow strips resulting from the SeqA region DETECTR assay.

The activation of the Cas12a protein through cleavage of target amplicons triggers the trans-cleavage of ssDNA probes containing FITC and biotin, enabling the visualization of TR4 genomic via LFA (**Fig. 2**). All crRNAs from the SCAR region showed a strong test (T) line (**Fig. 2C**). Intriguingly, for the SeqA region, a strong test (T) line was observed with crRNA2(I), even though exhibited lower cleavage activity compared to crRNA1(I) (**Fig. 2E,F**). This indicates a lack of correlation between crRNA cleavage activity and collateral probe cleavage activity. Considering this, crRNAs (crRNA1(S), crRNA2(S), crRNA3 (S)) from SCAR marker and crRNA2(I) from SeqA region) exhibiting both CRISPR cleavage activity and collateral cleavage activity of the probe were found suitable.

### Identification of suitable primers for RPA-DETECTR assay

In our preliminary studies, we tried amplifying the SCAR and SeqA regions using published primer sets (Li et al. 2013a; Ordóñez et al. 2019), however could not succeed. Therefore, 13 new primer combinations for the RPA amplification of the targets were designed to evaluate and optimize the DETECTR assay specificity (**Table 1**). To determine the optimal primer combinations for specificity in RPA assays, we conducted primer screening using genomic DNAs from Foc R1 and Foc TR4 as templates in RPA reactions. Following the screening process, we identified the AD, AF (from SeqA region), and GI (SCAR marker region) primer combinations (**Table 1**) as the most suitable for Foc TR4 detection (See Additional file 1: **Fig. S2**), exhibiting minimal background signal upon amplification and no amplification when Foc R1 genomic DNA was used as a template.

### Specificity of RPA-DETECTR system

In order to further analyses the specificity of the TR4-specific RPA primers (AD, AF, and GI primer combinations), we included a nonpathogenic Fo from garlic and Foc R1, as well as non-infected banana samples, as negative controls in the RPA assay. Remarkably, AF primer combination yielded highly specific amplification in the Fo and Foc R1, as well as in the non-infected banana samples compared to the AD and GI primer sets (**Fig. 3A**) and therefore AF primer targeting SeqA region was chosen for subsequent RPA-DETECTR assays.

**Fig. 3.**
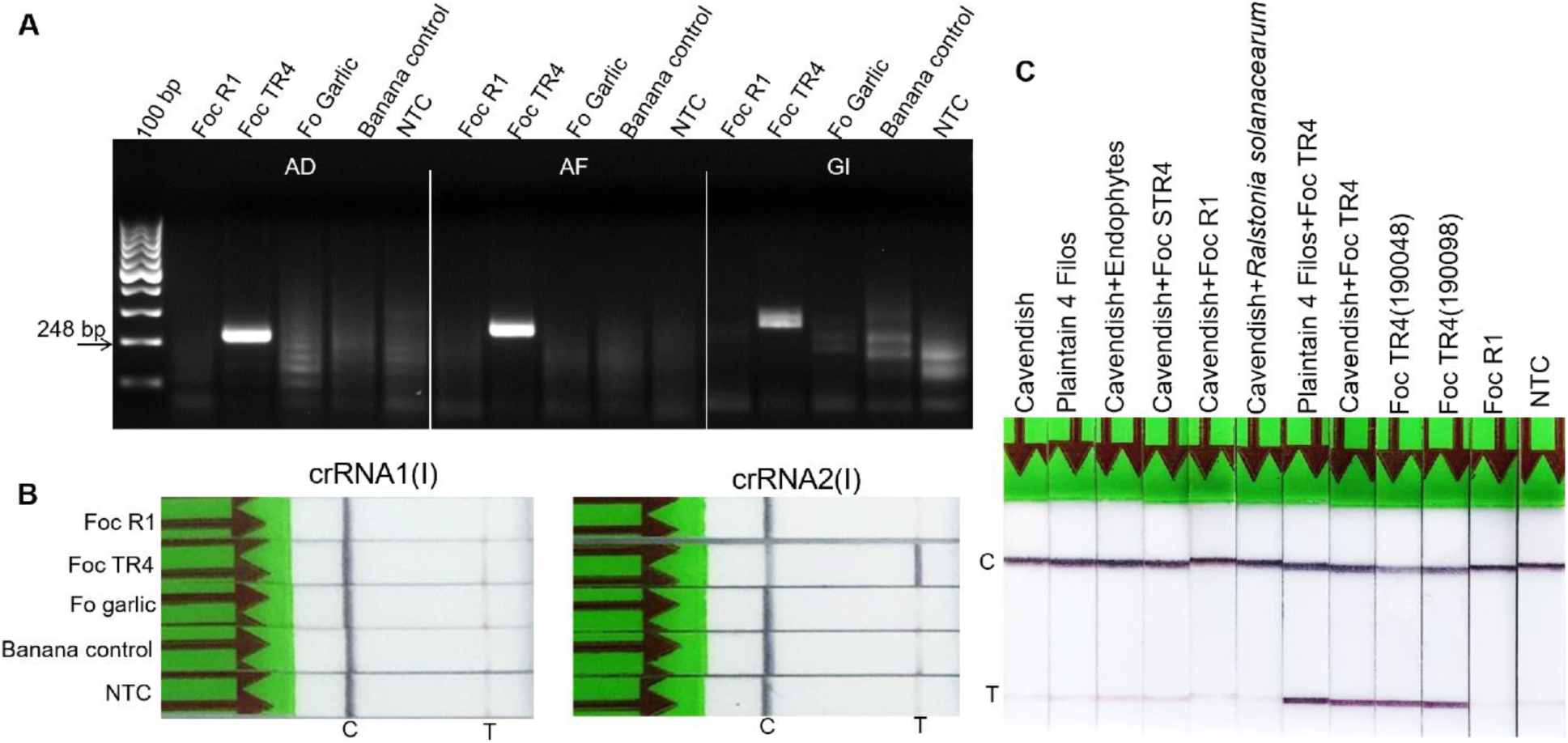
Recombinase polymerase amplification (RPA)-based DETECTR assays for Foc TR4 detection. **A**, Screening of primers for target amplification from the SeqA sequence using AD and AF primer combinations and the SCAR marker region using GI primer combinations. After RPA amplification and column purification, the reaction products were loaded onto 2.0% TBE agarose gel for analysis. **B**, Validation of crRNAs for the DETECTR assay. **C**, Assessment of the specificity of the Foc TR4 SeqA-specific DETECTR assay. NTC, no template control.

For the DETECTR assay, crRNA1(I) and crRNA2(I) from the SeqA region were evaluated. As expected, crRNA1(I) did not produce a positive T line signal in the LFA with Foc TR4 samples, whereas a positive signal was observed for Foc TR4 using crRNA2(I) (**Fig. 3B**). Furthermore, no obvious test line signal was observed with Fo garlic, Foc R1, and control banana samples, revealing no cross-reactivity of crRNA2(I) with Fo garlic, Foc R1, or banana genomic DNA. These findings indicate the specificity and efficacy of crRNA2(I) in the RPA-DETECTR assays. Additionally, these results were consistent with the PCR-based DETECTR assay using TR4-specific primers (**Fig. 2**).

### Foc TR4 detection using DETECTR

To validate the robustness and specificity of the DETECTR assay, a diverse panel of biological samples containing genomic DNA from different banana genotypes and/or samples spiked with genomic DNA from pathogens and endophytes were used, including Cavendish, Plantain 4 Filos, Cavendish + *R. solanacearum*, Plantain 4 Filos + Foc TR4, Cavendish + Foc TR4, Cavendish+STR4 as well as other Foc TR4 strains (190048 and 190098). The assay demonstrated high specificity, as the DETECTR signal was observed only in banana samples containing TR4 genomic DNA (**Fig. 3C**) while no signal was observed in samples including Cavendish, Plantain 4 Filos, and Cavendish + *R. solanacearum* and Cavendish+STR4. Thus, validation with a diverse sample panel confirmed the robustness and specificity of the DETECTR assay, detecting Foc TR4 only in banana samples with TR4 genomic DNA, with no false positive signal observed with other samples.

### Sensitivity analysis

To test the sensitivity of the assay, genomic DNA from Foc TR4 at various concentrations, ranging from 5.0 ng to 0.005×10^−3^ ng, was used (**Fig. 4**). At lower DNA concentrations, faint bands were visible on agarose gel (**Fig. 4A**). The DETECTR assay demonstrated high sensitivity, with Foc TR4-specific signal being visible at DNA concentrations as low as 5.0×10^−3^ ng (**Fig. 4A-C**), successfully detecting Foc TR4. We also analysed the sensitivity using infected banana samples (three weeks after inoculation) with genomic DNA ranging from 5.0 ng to 0.3125 ng/reaction. In the case of three-week-old, infected banana plant samples, Foc TR4 can be detected using 1.25 ng/reaction (**Fig. 4D**).

**Fig. 4.**
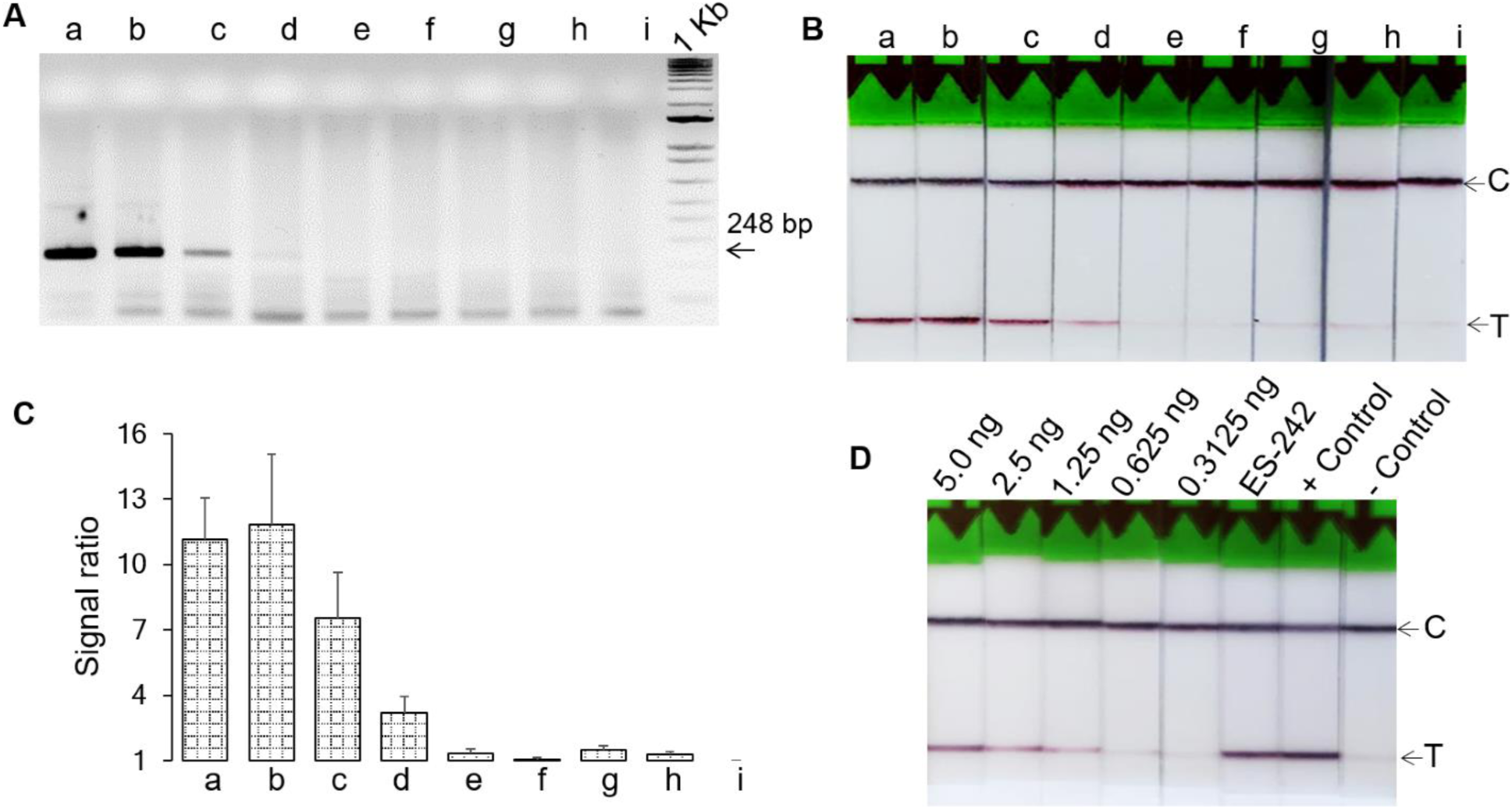
Sensitivity of Foc TR4 detection SeqA-specific detection assays. **A**, RPA amplification of Foc TR4 target region. 20 µl reaction products were column purified and visualized on 2.0% TBE agarose gel. **B**, Assessment of the specificity of Foc DETECTR assay. **C**, Normalized signal intensities of DETECTR assay. The error bars indicate the standard error of four independent experimental replicates. **D**, Sensitivity of Foc TR4 detection using Foc TR4 infected banana sample. a, b, c, d, e, f, and g correspond to 5.0, 0.5, 0.05, 5.0×10^−3^, 0.5×10^−3^, 0.05×10^−3^, 0.005×10^−3^ ng/reaction of Foc TR4 genomic DNA, and h and i corresponds to no template controls (NTC).

To determine if the DETECTR assay exhibits higher sensitivity compared to PCR, qPCR, or ddPCR, the detection levels of Foc TR4 using each method were compared (**Fig. 5 A-D**). While conventional PCR could reliably detect the pathogen only at 0.05 ng/reaction (**Fig. 5C**), both qPCR and ddPCR demonstrated high sensitivity, detecting Foc TR4 down to 0.5-0.005×10^−3^ ng/reaction (**Fig. 5A, B, D**). However, as the template concentration decreased in qPCR, the formation of primer dimers led to misleading results at lower template concentrations. In contrast, while 5.0×10^−3^ ng/reaction was sufficient to detect the Foc TR4 pathogen in a 25-min RPA-DETECTR reaction it offered high sensitivity and rapidity in Foc TR4 detection particularly due to its quick amplification process, surpassing conventional PCR and addressing the limitations observed in qPCR.

**Fig. 5.**
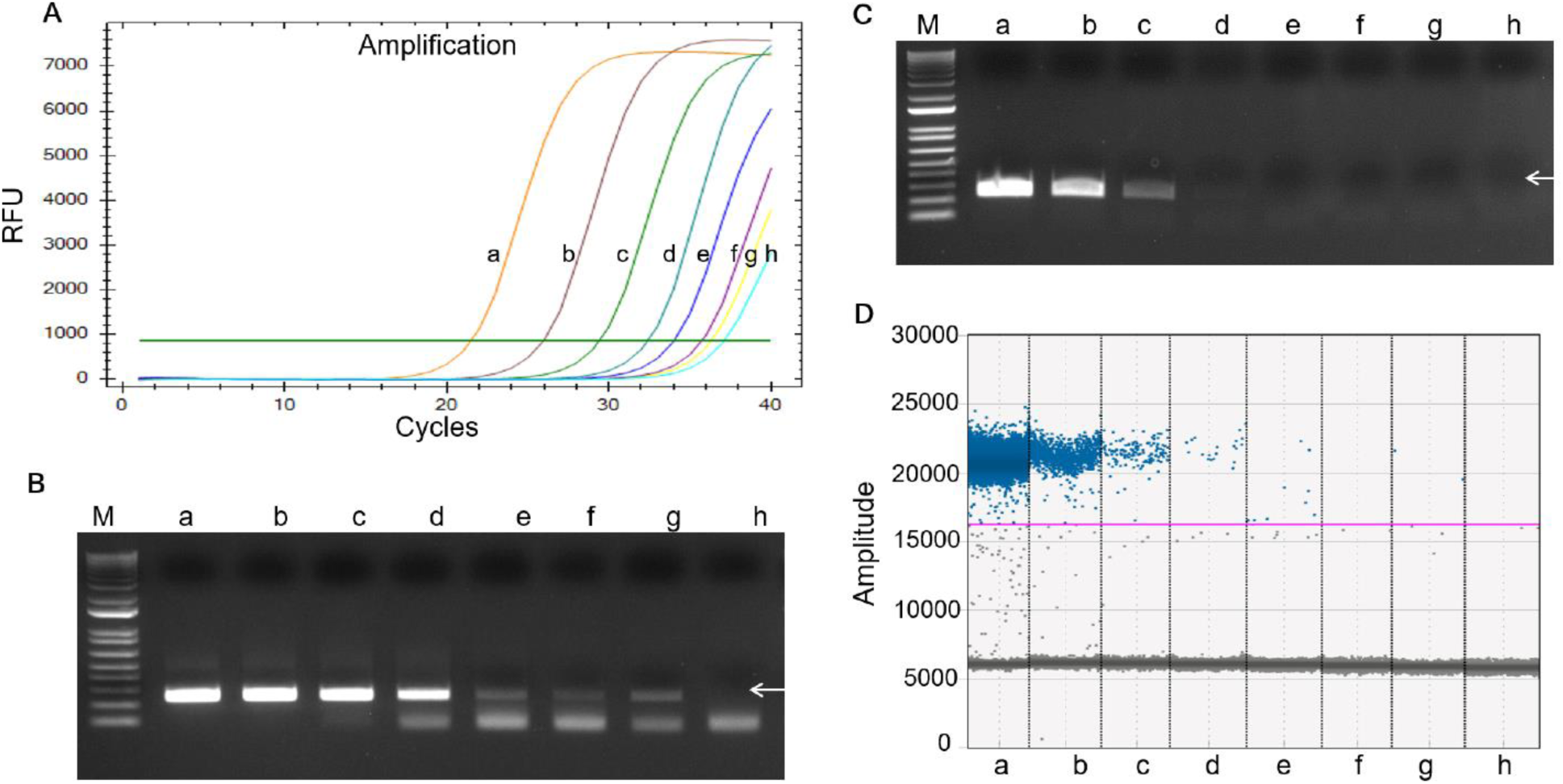
Comparative sensitivity analysis of Foc TR4 detection using SeqA-specific AF primers through PCR, qPCR and ddPCR. **A**, qPCR sensitivity analysis. **B**, qPCR amplification products of Foc TR4 target region visualized through gel electrophoresis. **C**, Sensitivity assessment of Foc TR4 using PCR assay. **D**, Sensitivity assessment of Foc TR4 using ddPCR assay. a, b, c, d, e, f, and g correspond to 5.0, 0.5, 0.05, 5.0×10^−3^, 0.5×10^−3^, 0.05×10^−3^, 0.005×10^−3^ ng/reaction genomic DNA of Foc TR4, and h corresponds to the no template control (NTC). The arrowheads in the gel correspond to 248 bp.

The robustness of the DETECTR assay was further validated using Foc TR4 inoculated Cavendish banana plants showing low to moderate cross sectional xylem tissue browning three weeks after inoculation (**Fig. 6A**). PCR analysis following 36 cycles of amplification using 4.0 ng of template DNA per reaction showed presence of Foc TR4 infection in these tissues (**Fig. 6B**), with PCR band intensities correlating with the infection levels of the samples. In pseudostems with weak or no visual symptoms, PCR showed weak signals. The DETECTR assay produced distinct signals in the root, corm, and collar tissues of even in symptomless plants distinguishable from the background signal of non-template control (NTC) samples (**Fig. 6C**), demonstrating its effectiveness for diagnosing banana Fusarium wilt as early as three weeks post-inoculation, using as little as 2.0 ng of genomic DNA per reaction. Overall, this study suggests the efficient amplification of the Foc TR4-specific genomic target through RPA and subsequent detection through the DETECTR assay.

**Fig. 6.**
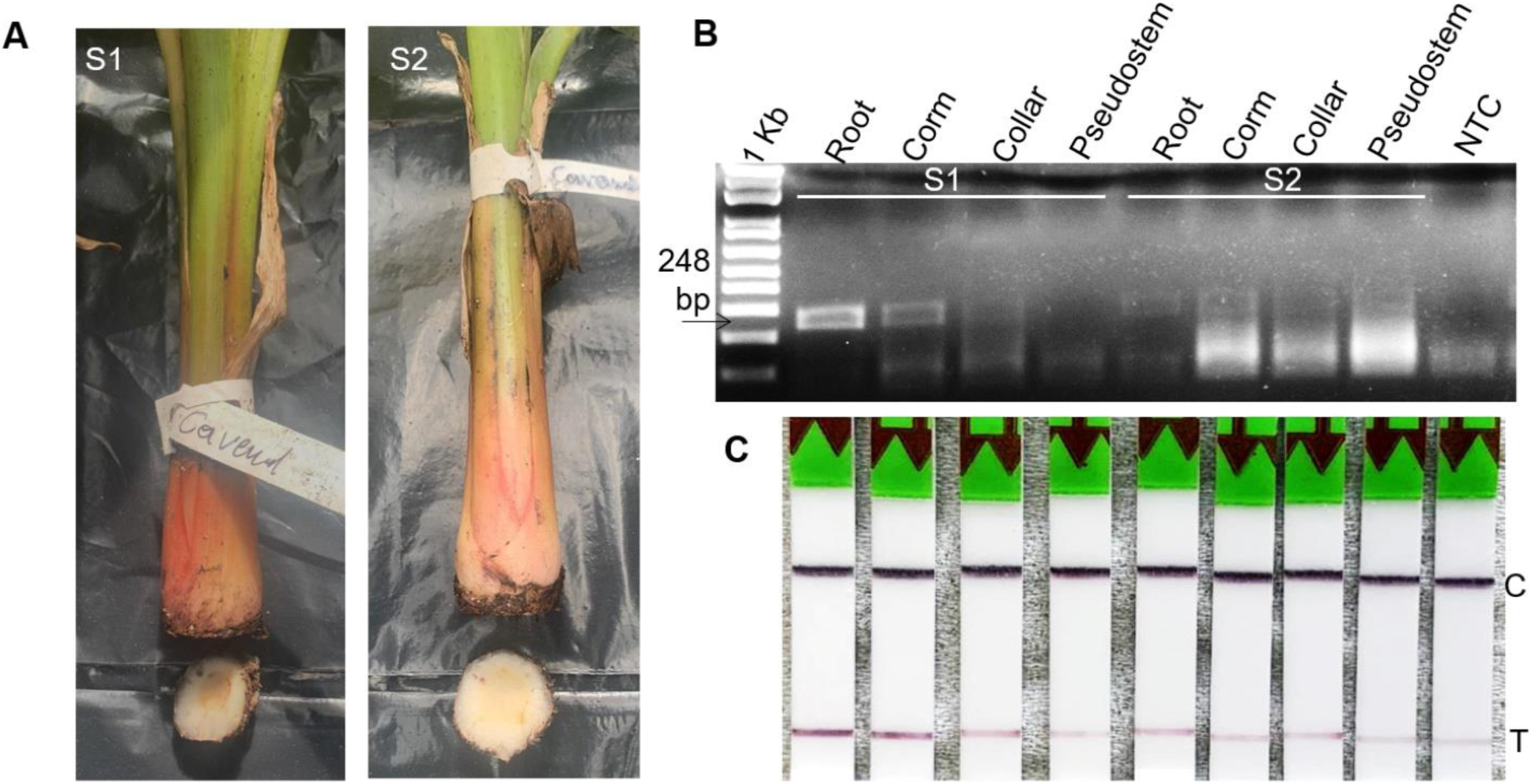
Detection of Foc TR4 infected samples using the DETECTR assay. **A**, Visual symptoms of a Foc TR4 infected Cavendish banana plant samples (S1: symptomatic, S2 asymptomatic) three weeks post-inoculation. **B**, PCR detection of Foc TR4 using AF primers (corresponding to 36 cycles of amplification and 4.0 ng of per PCR reaction). **C**, DETECTR assay for Foc TR4 detection three weeks post-inoculation. NTC, no template control.

## Discussion

The emergence of CRISPR/Cas nucleic acid detection, particularly the DETECTR assay marks a significant advancement in diagnostic technology, leveraging the transformative capabilities of CRISPR/Cas for detecting specific targets through the trans-cleavage of single-stranded DNA reporters (Chen et al. 2018; Yamano et al. 2016). The DETECTR assay, known for its rapidity, portability, ultra-sensitivity, and cost-effectiveness, has emerged as a viable method for detecting pathogens (Broughton et al. 2020). Recent research harnessing the collateral cleavage activity of CRISPR/Cas system has primarily focused on detecting nucleic acids from human pathogens (Broughton et al. 2020), but there has been limited attention given to disease-causing plant pathogens.

Banana Fusarium wilt causes devastating crop damage and is included in the list of quarantine pests in various countries (EFSA PLH Panel et al. 2022; IPPC 2023). To increase disease diagnosis efficacy, expedite customs clearance, and prevent cross-border dissemination, a rapid and accurate detection approach is imperative. This study reports the first successfully developed RPA-DETECTR assay for the highly sensitive and specific detection of banana wilt pathogen Foc TR4 and its validation across several isolates. To customize the DETECTR method for robust Foc TR4 detection, specific crRNA spacer sequences and RPA primers were designed and evaluated for specificity and sensitivity. Choosing appropriate and specific target crRNA sites is crucial to enhance the viability and effectiveness of crRNAs for DETECTR assay. In our study, at least three crRNAs were used for the RPA-DETECTR assay for each target region, depending on the presence of the TTTV PAM site. Our focus was on two regions previously reported as specific to Foc TR4 (Li et al. 2013a; Ordóñez et al. 2019). Our findings revealed varying cleavage activities among the designed crRNAs. High cleavage activity was observed in all three crRNAs from the SCAR marker region (Li et al. 2013a), while only crRNA1(I) from the SeqA region showed relatively high cis-cleavage activity. However, crRNA1(I) did not exhibit trans-cleavage activity, as indicated by the absence of a T line signal in the LFA. Therefore, we opted to utilize crRNA2(I) from the SeqA region (Ordóñez et al. 2019) since it demonstrated trans-cleavage activity, despite having relatively weak cis-cleavage activity. A previous study has demonstrated varying levels of trans-cleavage activities among different crRNAs, with some showing little to no trans-cleavage activity (Weng et al. 2023). Therefore, designing and testing of multiple crRNAs for each target region is important to identifying crRNAs with both cis– and trans-nuclease activity for detection purposes.

Pre-amplification was deemed necessary for the DETECTR assay. In comparison to PCR, RPA proves more suitable for the early detection of Foc TR4 due to its lower requirements for DNA contents and simpler reaction conditions (Piepenburg et al. 2006). Due to nonspecific amplification observed with SCAR marker region-specific primers in the RPA assay, this region was excluded from the DETECTR assay. Utilizing isothermal amplification of the DETECTR target regions, we confirmed the specificity and efficacy of Foc TR4-specific primers designed from the SeqA sequence. The DETECTR Foc TR4 assay demonstrated high specificity for Foc TR4, and non-target species, such as *R. solanacearum* (a bacterial pathogen causing similar wilting symptoms), *Fusarium oxysporum*, Foc R1, and other endophytes, did not produce any signals indicating the high specificity of the assays. The sensitivity is notable, detecting as little as 0.005 ng of pure Foc TR4 DNA per reaction, with excellent reproducibility and amplification efficiency, even in the presence of background DNA. In contrast, conventional PCR reliably identified the pathogen at 0.05 ng/reaction. In our studies, qPCR and ddPCR showed higher sensitivity levels of Foc TR4 detection. However, we observed the formation of primer dimers as the template concentration decreased, leading to potentially misleading results at lower template concentrations down to 5.0×10^−3^ ng/reaction. While the DETETCR assay sensitivity is lower than the present and previously reported sensitivity of ddPCR (using 2.5×10^−3^ ng/reaction) (Lovera et al. 2024), and qPCR (using 1.0×10^−3^ ng/reaction) (Aguayo et al. 2017) assays, a 25-min RPA reaction showcased the sensitivity, aligning with the sensitivity levels of other Cas12a nucleases utilized for diagnostic purposes (Wheatley et al. 2021; Teng et al. 2019; Chen et al. 2018). Extending the RPA duration to 30 min could potentially enhance detection sensitivity further. Furthermore, the DETECTR assay effectively mitigates false-positive signals originating from nonspecific amplification products through crRNA recognition (Huang et al. 2023).

In terms of throughput, the RPA-DETECTR assays enabled faster detection of pathogens than other molecular diagnostics, with TR4 diagnosis in test samples completed within 60 min with minimal nucleic acid content (de Dieu Habimana et al. 2022). Consequently, serving as a ready-to-use POC tool for routine Foc TR4 diagnosis, supporting the implementation of cordoning and quarantine strategies and provide timely support in combating plant diseases in remote and limited-resource settings. In conclusion, our study for the first time report RPA-DETECTR assays for the detection of banana Foc TR4 with high specificity, PCR amplification-free detection, rapidity, and enhanced sensitivity.

The RPA-DETECTR assay demonstrated its capability to identify Foc TR4 within a brief 60-min timeframe successfully detecting infection in the plant tissues as well as a diverse panel specifically well suited for POC applications. This study lays the groundwork for potential enhancements that could result in straightforward and efficient field tests for early Foc TR4 detection.

## Declarations

### Ethics approval and consent to participate

Not applicable.

## Consent for publication

All the authors approved the manuscript for publication.

## Availability of data and materials

All the data generated or analysed during this research are included in this published article and supplementary information files.

## Authors’ contributions

PBM and JV conceptualized the study. PBM, JV, JJ-C, and MM were involved in the planning and development of the assays. JV, HM, and AA participated in disease inoculation and sample preparation. JV, MS-S, CZ, and ICCB supported the validation of the pathogen samples and panel. JV and PBM prepared the manuscript. All the authors contributed to the manuscript.

## Acknowledgments

We would like to express our sincere gratitude to Dr. Viljoen Altus for kindly providing the Foc TR4 and Foc R1 strains from Africa. This research contributes to the objectives of the CRP D23033 An Integrative Approach to Enhance Disease Resistance Against Fusarium Wilt (Foc TR4) in Banana – Phase II and Peaceful Use Initiative (PUI) project on “Enhancing climate change adaptation and disease resilience in banana–coffee cropping systems in East Africa’.

---

**Table S1.**
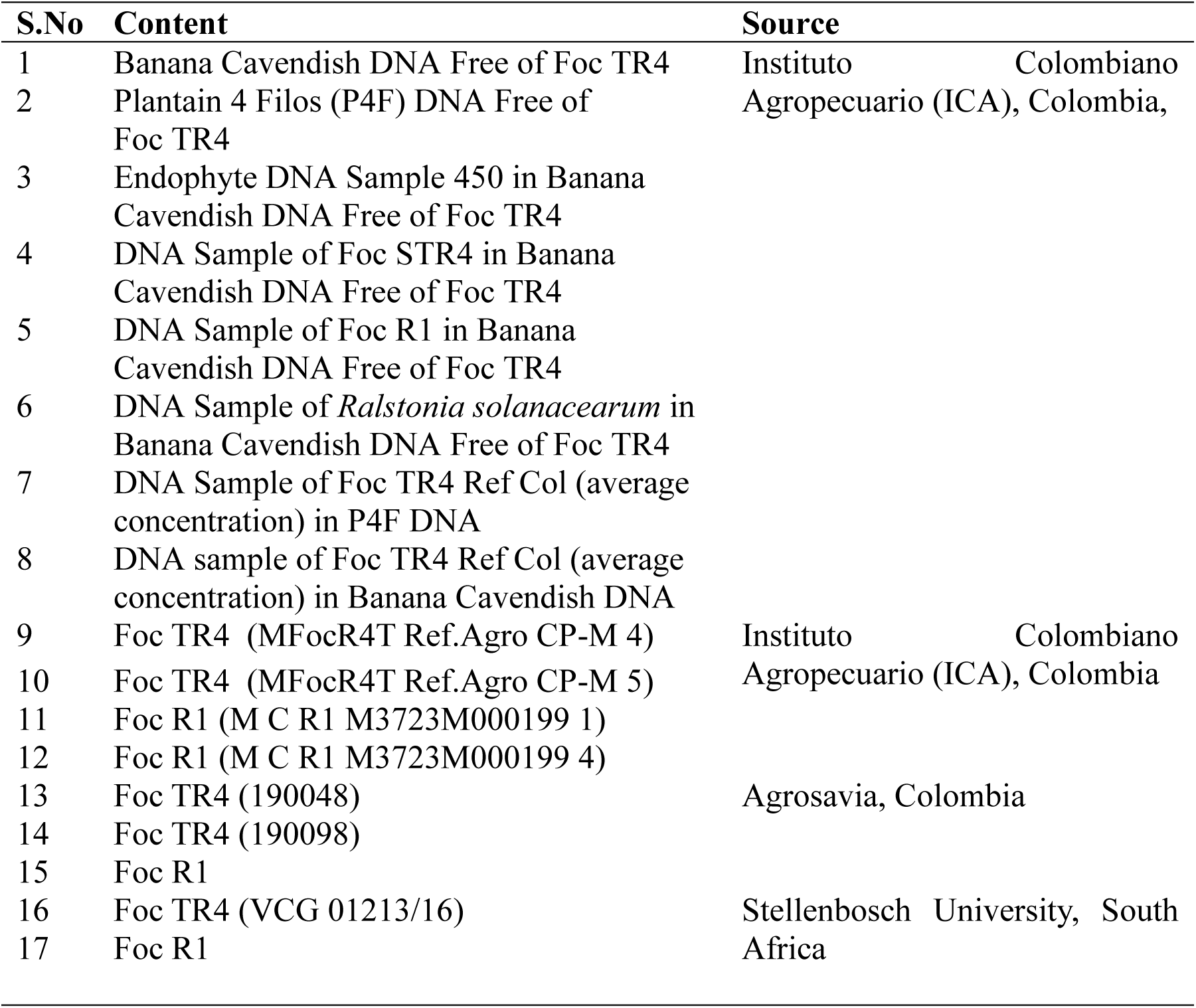
Details of detection panels and *Fusarium oxysporum* strains used in the present study.

**Fig. S1.**
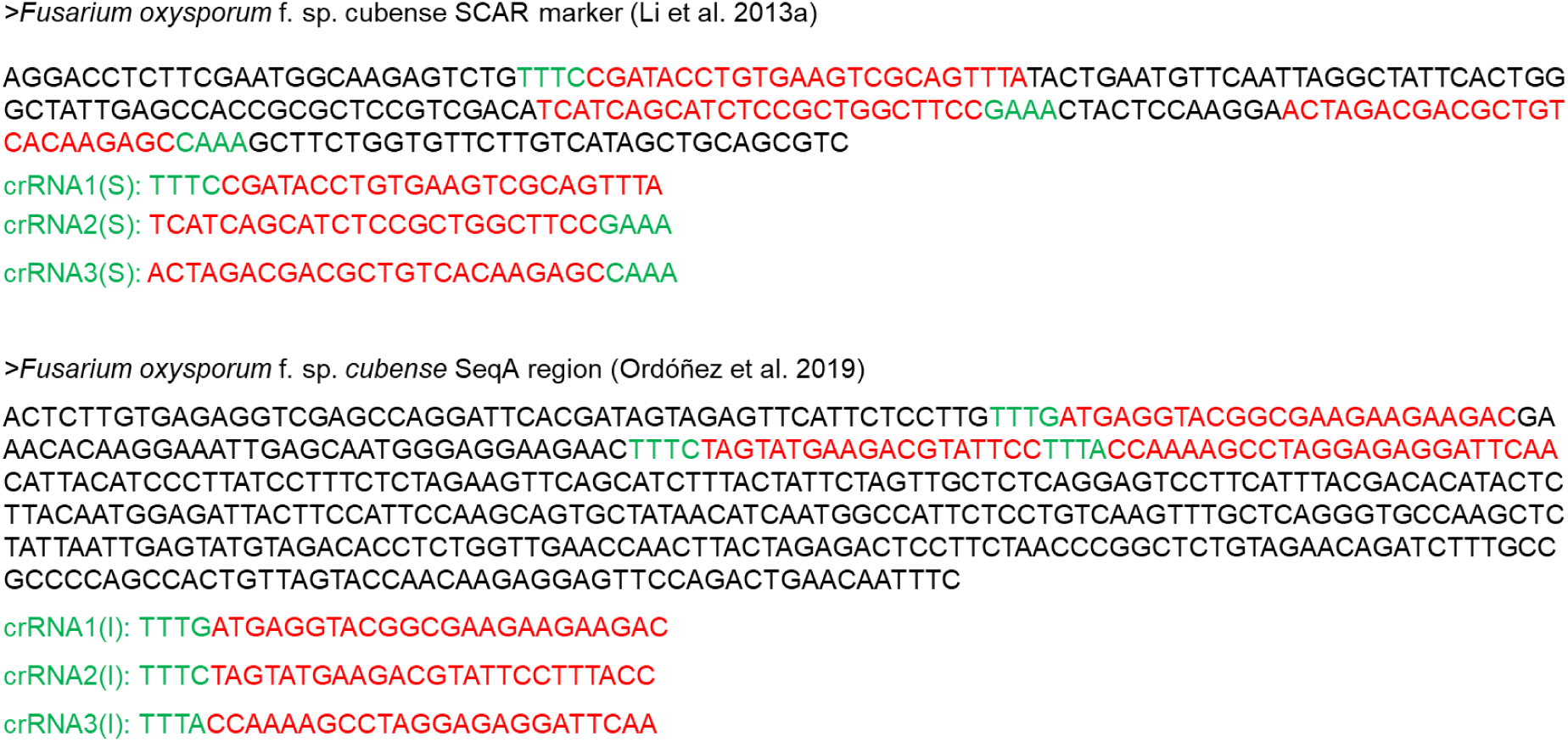
Foc TR4 specific SCAR marker and SeqA regions and crRNA sites. crRNA sequences are in red font. PAM sites are in green font.

**Fig. S2.**
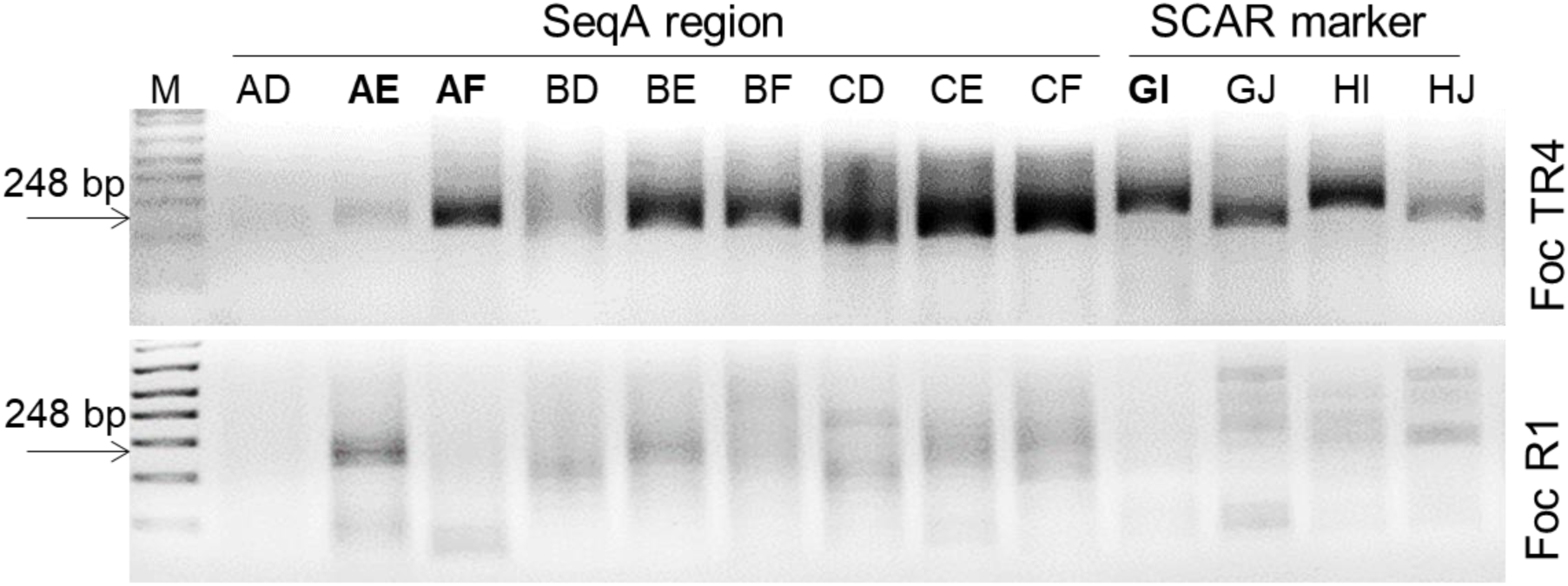
Recombinase polymerase amplification (RPA) primer evaluation. RPA testing of primer combinations for ITS target amplification. RPA products were column purified and visualized using 2.0% TAE agarose gel electrophoresis. An efficient amplification with three RPA primer combinations (AD, AF and GI) were selected for subsequent RPA assays.

