## Additional file 1 for "Rapid and Sensitive Detection of *Fusarium oxysporum* f. sp. *cubense* Tropical Race 4 Using a RPA-DETECTR Assay"

**Table S1.** Details of detection panels and *Fusarium oxysporum* strains used in the present study.

| S.No | Content | Source |
|---|---|---|
| 1 | Banana Cavendish DNA Free of Foc TR4 | Instituto Colombiano |
| 2 | Plantain 4 Filos (P4F) DNA Free of Foc TR4 | Agropecuario (ICA), Colombia, |
| 3 | Endophyte DNA Sample 450 in Banana Cavendish DNA Free of Foc TR4 | |
| 4 | DNA Sample of Foc STR4 in Banana Cavendish DNA Free of Foc TR4 | |
| 5 | DNA Sample of Foc R1 in Banana Cavendish DNA Free of Foc TR4 | |
| 6 | DNA Sample of *Ralstonia solanacearum* in Banana Cavendish DNA Free of Foc TR4 | |
| 7 | DNA Sample of Foc TR4 Ref Col (average concentration) in P4F DNA | |
| 8 | DNA sample of Foc TR4 Ref Col (average concentration) in Banana Cavendish DNA | |
| 9 | Foc TR4 (MFocR4T Ref.Agro CP-M 4) | Instituto Colombiano |
| 10 | Foc TR4 (MFocR4T Ref.Agro CP-M 5) | Agropecuario (ICA), Colombia |
| 11 | Foc R1 (M C R1 M3723M000199 1) | |
| 12 | Foc R1 (M C R1 M3723M000199 4) | |
| 13 | Foc TR4 (190048) | Agrosavia, Colombia |
| 14 | Foc TR4 (190098) | |
| 15 | Foc R1 | |
| 16 | Foc TR4 (VCG 01213/16) | Stellenbosch University, South |
| 17 | Foc R1 | Africa |

>*Fusarium oxysporum* f. sp. cubense SCAR marker (Li et al. 2013a)

AGGACCTCTTCGAATGGCAAGAGTCTGTTTCCGATACCTGTGAAGTCGCAGTTTATACTGAATGTTCAATTAGGCTATTCACTGG
GCTATTGAGCCACCGCGCTCCGTCGACATCATCAGCATCTCCGCTGGCTTCCGAAACTACTCCAAGGAACTAGACGACGCTGT
CACAAGAGCCAAAGCTTCTGGTGTTCTTGTCATAGCTGCAGCGTC

crRNA1(S): TTTCCGATACCTGTGAAGTCGCAGTTTA

crRNA2(S): TCATCAGCATCTCCGCTGGCTTCCGAAA

crRNA3(S): ACTAGACGACGCTGTCACAAGAGCCAAA

>*Fusarium oxysporum* f. sp. *cubense* SeqA region (Ordóñez et al. 2019)

ACTCTTGTGAGAGGTCGAGCCAGGATTCACGATAGTAGAGTTCATTCTCCTTGTTTGATGAGGTACGGCGAAGAAGAAGACGA
AACACAAGGAAATTGAGCAATGGGAGGAAGAACTTTCTAGTATGAAGACGTATTCCTTTACCAAAAGCCTAGGAGAGGATTCAA
CATTACATCCCTTATCCTTTCTCTAGAAGTTCAGCATCTTTACTATTCTAGTTGCTCTCAGGAGTCCTTCATTTACGACACATACTC
TTACAATGGAGATTACTTCCATTCCAAGCAGTGCTATAACATCAATGGCCATTCTCCTGTCAAGTTTGCTCAGGGTGCCAAGCTC
TATTAATTGAGTATGTAGACACCTCTGGTTGAACCAACTTACTAGAGACTCCTTCTAACCCGGCTCTGTAGAACAGATCTTTGCC
GCCCCAGCCACTGTTAGTACCAACAAGAGGAGTTCCAGACTGAACAATTTC

crRNA1(I): TTTGATGAGGTACGGCGAAGAAGAAGAC

crRNA2(I): TTTCTAGTATGAAGACGTATTCCTTTACC

crRNA3(I): TTTACCAAAAGCCTAGGAGAGGATTCAA

**Fig. S1**. Foc TR4 specific SCAR marker and SeqA regions and crRNA sites. crRNA sequences are in red font. PAM sites are in green font.

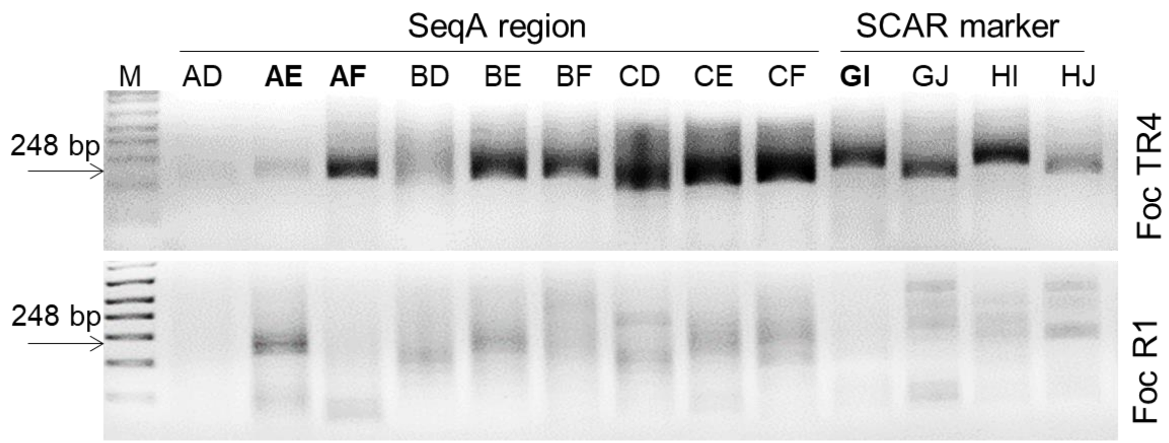


**Fig. S2**. Recombinase polymerase amplification (RPA) primer evaluation. RPA testing of primer combinations for ITS target amplification. RPA products were column purified and visualized using 2.0% TAE agarose gel electrophoresis. An efficient amplification with three RPA primer combinations (AD, AF and GI) were selected for subsequent RPA assays.
